# HSP90.6 is involved in grain filling via carbon and nitrogen metabolism in maize

**DOI:** 10.1101/2022.04.27.489727

**Authors:** Jianghai Xu, Zhijia Yang, Xiaohong Fei, Meiling Zhang, Yang Cui, Xiangbo Zhang, Kaiwen Tan, Lizhu E, Haiming Zhao, Jinsheng Lai, Qian Zhao, Weibin Song

## Abstract

Carbon and nitrogen are the two most abundant nutrients in all living things, and their metabolism maintains normal plant growth. However, the molecular mechanism underlying carbon and nitrogen metabolism remains largely unknown. Here, we found that HSP90.6 is involved in the metabolism of carbon and nitrogen. We performed gene cloning and functional characterization of a maize EMS mutant *ehsp90.6*, whose kernels were small. *HSP90.6* encodes heat shock protein 90.6, which has a single-amino acid mutation within its HATPase_c domain. Transcriptome profiling showed that the expression of amino acid biosynthesis- and carbon metabolism-related genes was significantly downregulated in *hsp90.6*. HSP90.6 is involved in the 26S proteasome degradation pathway, which affects nitrogen recycling to regulate amino acid synthesis; this occurs by interactions between HSP90.6 and the 26S proteasome subunits RPN6 and PBD2 (PRC2). The loss of HSP90.6 significantly reduced the activity of the 26S proteasome, resulting in the accumulation of ubiquitinated proteins and defects in nitrogen recycling. Moreover, HSP90.6 interacted with the 14-3-3 protein GF14-6 to participate in carbon metabolism. Together, these findings revealed that HSP90.6 regulates nutrient metabolism in maize seeds by affecting 26S proteasome-mediated nitrogen recycling and GF14-6-mediated carbon metabolism.

**One sentence summary:** HSP90.6 is involved in nutrient metabolism via 26S proteasome-mediated protein degradation to promote nitrogen recycling and GF14-6 protein-mediated carbon metabolism.

The author responsible for the distribution of materials integral to the findings presented in this article in accordance with the policy described in the Instructions for Authors (https://academic.oup.com/plcell/pages/General-Instructions) is Weibin Song.

**Highlights:**

- HATPase_c is necessary for HSP90.6 to regulate maize kernel development.
- HSP90.6 is involved in nitrogen recycling via the 26S proteasome degradation pathway.
- HSP90.6 interacts with the 14-3-3 protein GF14-6 to affect carbon metabolism.

**IN A NUTSHELL:** *Background:* Seeds are the main harvested organs of maize. Understanding the regulatory mechanism of grain filling is helpful to cultivate high-quality and high-yield maize. In the past few years, the regulatory network of grain filling has been explored through multiple means, including transcriptomic, proteomic and functional genomic techniques. Many genes that control grain filling through different mechanisms have been cloned, such as *CTLP1 (Choline Transporter-like Protein 1), OS1* (*Opaque Endosperm and Small Germ 1*), and *MN6* (*Miniature Seed6*). To identify new genes involved in maize grain filling, ethyl methanesulfonate (EMS) was used to induce mutations, and the *ehsp90.6* mutant, which exhibited abnormal kernel development, was isolated by bulked segregant analysis RNA sequencing (BSR).

*Question:* Why does the single-amino acid mutation of HSP90.6 affect grain size, and how does the loss of HSP90.6 affect grain filling?

*Findings:* A single-amino acid mutant (*ehsp90.6*) and knockout mutant (*hsp90.6*) were obtained. We found that HSP90-6 was involved in the regulation of maize grain filling. A single-single amino acid mutation in the HATPase_c domain reduced the ATPase activity of HSP90.6, resulting in smaller grains. The functional loss of HSP90.6 resulted in the expression of amino acid biosynthesis- and carbon metabolism-related genes being significantly downregulated in *hsp90.6*. We indicated that HSP90.6 is involved in the 26S proteasome degradation pathway, which affects nitrogen recycling to regulate amino acid synthesis by interacting with the 26S proteasome subunits RPN6 and PBD2 (PRC2). Moreover, HSP90.6 was found to interact with the 14-3-3 protein GF14-6 to participate in carbon metabolism.

*Next steps:* To further verify that the interaction between HSP90.6 and 26S proteasome subunits and GF14-6 affects grain filling, knockout validation of RPN6, PBD2 (PRC2) and GF14-6 will be performed. In addition, since GF14-6 interacts with the phosphorylated proteins, we will determine the phosphorylation site of HSP90.6. Due to the important role of HSP90 family proteins in plant development, there are other regulatory pathways that need to be explored.

## Introduction

Carbon (C) and nitrogen (N) are two essential macronutrients required for plant development, and these nutrients maintain plant growth and accumulate within biomass (Gao et al., 2020). C/N metabolism is inseparable from grain development, and the grains are the main organs for increasing production. Understanding the C and N regulatory mechanisms is helpful for cultivating high-quality and high-yield crops. In the past few years, the regulatory network of kernel development has been explored and many genes that control seed development through different mechanisms have been cloned, such as *CTLP1 (Choline Transporter-like Protein 1), OS1* (*Opaque Endosperm and Small Germ 1*), and *MN6* (*Miniature Seed6*) (Hu et al., 2021; Song et al., 2019; Yi et al., 2021). However, the regulatory network involving C/N metabolism and grain development is still unclear. It has been reported that some heat shock proteins are involved in nitrogen and amino acid metabolism in the endosperm (Ishimaru et al., 2019), but it remains unclear whether HSP90 is also involved in this process.

Heat shock protein 90 (HSP90) is highly conserved chaperone protein subfamily (Schopf et al., 2017), belonging to the GHKL (Gyrases, HSP90, histidine kinase and MutL) superfamily of ATPases (Dutta and Inouye, 2000). HSP90 contains three domains: an N-terminal ATPase domain responsible for hydrolyzing ATP to release energy, an intermediate domain responsible for client protein binding, and a C-terminal dimerization domain (Pearl and Prodromou, 2006; Prodromou, 2016). HSP90 plays important roles in cell signal transduction (Jackson et al., 2004; Picard et al., 1990; Pratt, 1997), the cell cycle and maintaining protein integrity and homeostasis (McClellan et al., 2005; Taipale et al., 2010; Young et al., 2003). HSP90 was named according to the 90 kDa group of proteins in which it belongs and whose expression is upregulated in stressed cells. HSP90s have also been found to be important parts of the cell development process under normal physiological conditions, according to in-depth research (Krishna and Gloor, 2001; Rutherford and Lindquist, 1998). Loss of function of HSP90 affects the embryogenesis, seed germination and pollen development (Cha et al., 2013; Feng et al., 2013; Luo et al., 2019; Prasinos, 2005).

Generally, HSPs function as molecular chaperones by interacting with multiple types of proteins (Hoter et al., 2018; Prodromou, 2016). HSP90 and HSP70 interact with the 26S proteasome in yeast and participate in the degradation of ubiquitinated proteins (Park et al., 2007; Verma et al., 2000). In addition, ScHSP110 interacts with the 19S regulatory particles (RPs) of the 26S proteasome and then recruits the substrate-HSP70 complex to participate in protein degradation via the 26S proteasome (Kandasamy and Andréasson, 2018). The 26S proteasome is the key complex responsible for regulating protein degradation in eukaryotic cells and controlling many cellular processes, including the cell cycle, DNA replication and signal transduction (Bhattacharyya et al., 2014; Collins and Goldberg, 2017; Finley, 2009; Goldberg, 2007). The 26S proteasomal degradation pathway has been considered to be a recycler of proteins and thus a contributor to N homeostasis (Tornkvist et al., 2019). Under conditions of amino acid starvation, autophagy and proteasomal degradation are required to maintain sufficient amino acid levels to sustain protein synthesis (Elharar et al., 2014; Tornkvist et al., 2019). Proteasomal degradation exerts trophic effects through amino acid cycling (Suraweera et al., 2012). Research shows that proteasome inhibition can lead to fatal amino acid shortages. In yeast, mammals and *Drosophila*, the deleterious consequences of proteasome inhibition can be rescued by amino acid supplementation (Suraweera et al., 2012).

In addition to being molecular chaperones, HSPs can also perform regulatory functions through posttranslational modification, including phosphorylation, acetylation and methylation (Prodromou, 2016; Sima and Richter, 2018). Phosphorylation is the most common posttranslational modification involved in regulating the activity of most HSPs (Hornbeck et al., 2012; Muller et al., 2013; Nguyen et al., 2017). 14-3-3 proteins are highly conserved dimeric proteins that bind to Ser/Thr phosphorylated sites on substrate proteins (Madeira et al., 2015; Morrison, 2009). It has been reported that the 14-3-3 protein Bmh1 recruits phosphatase Glc7 by interacting with HSP70 to participate in the regulation of sugar metabolism in *Saccharomyces cerevisiae* (Hübscher et al., 2016; Chen et al., 2019). In *Arabidopsis* and maize, there are 13 and 12 14-3-3 proteins, 150 respectively, of which 7 are present in the seeds of both species. The 14-3-3 isoforms chi and epsilon in *Arabidopsis* (Swatek et al., 2011) and GF14-4 and GF14-6 in maize are significantly expressed in the seeds (Dou et al., 2015). Protein–protein interactions between substrate proteins show that most substrate proteins are involved in glycolysis, fatty acid synthesis and protein storage (Dou et al., 2015; Ma et al., 2016; Swatek et al., 2011). This is in line with the findings of previous studies based on several specific targets of 14-3-3s: 14-3-3s mainly regulate C/N metabolism in plants (Comparot et al., 2003; Diaz et al., 2011; Fu et al., 2000; Fulgosi et al., 2002). However, the specific underlying mechanism is still unclear.

Here, we cloned *HSP90.6* from maize. This gene is involved in the regulation of grain filling via carbon and nitrogen metabolism during maize kernel development. ATPase activity is affected because a single amino acid altered the HATPase_c domain sequence, which ultimately resulted in smaller kernels. We demonstrated that HSP90.6 regulates amino acid biosynthesis via the 26S proteasome degradation pathway, which affects nitrogen recycling. Furthermore, HSP90.6 was found to affect carbon metabolism by interacting with the 14-3-3 protein GF14-6 to participate in grain filling. This research revealed that HSP90.6 regulates nutrient metabolism during maize kernel development by 26S proteasome-mediated nitrogen recycling and GF14-6 mediated carbon metabolism.

## Results

### Cloning and identification of *HSP90.6*

To identify new genes involved in maize kernel development, ethyl methanesulfonate (EMS) was used to induce mutations, and a mutant with abnormal kernel development, *ehsp90.6*, was isolated by bulked segregant analysis RNA sequencing (BSR). A single-nucleotide mutation (G380A) was identified, which resulted in a single amino acid change from Arg to Gln (Fig. 1A, B and D; Table S1). Compared with wild-type (WT) mature seeds, mature seeds of *ehsp90.6* were smaller and slightly shrunken (Fig. 1A and B; Fig. S1, A-C). The mutant seeds could germinate normally, but the seedlings showed dwarfism (Fig. 1C; Fig. S1D and E). The segregation ratio of normal kernels and smaller kernels was approximately 3:1, indicating that the mutation in *ehsp90.6* is a single-gene recessive one. To further determine the candidate gene, three independent knockout plants were obtained, all of which were heterozygous mutants with an insertion or deletion at the target site leading to frameshift mutations (Fig. S2A). When T1-generation heterozygous mutants were selfed, some seeds on the mature ears showed obvious shrinkage, almost no filling, and could not germinate (Fig. S2B); the segregation ratios were close to 3:1 (246:74, 352:127, 298:112). The progeny resulting from the hybridization of plants with the different transformation plants also showed the same results (Fig. S2D).

**Figure 1.**
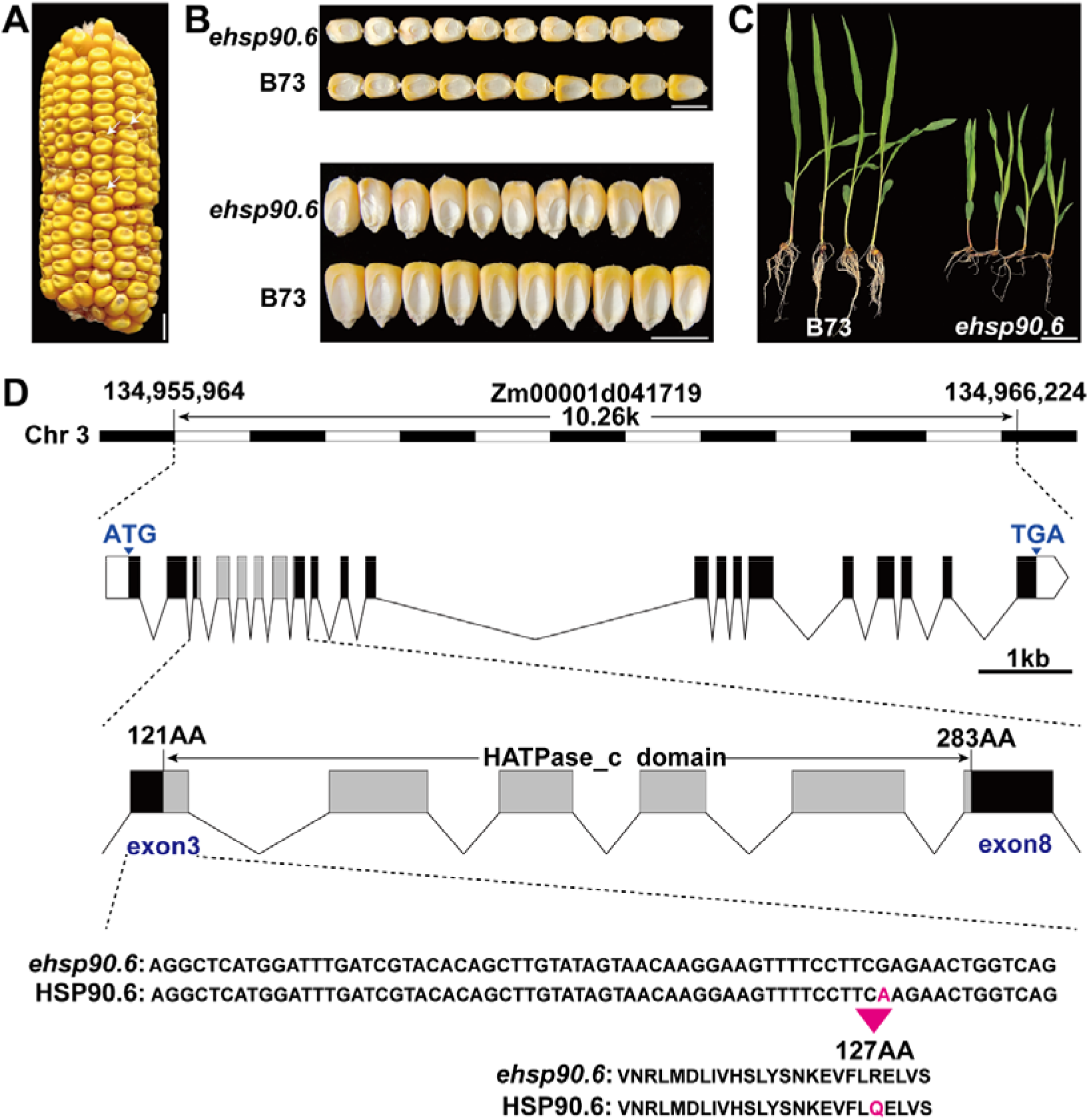
Cloning and phenotypic characteristics of *ehsp90.6* A, Ears of heterozygous *ehsp90.6* F2 mutants; the magenta arrow represents homozygous *ehsp90.6*. Scale bar =1 cm. B, Mature seeds of B73 and *ehsp90.6*. Scale bar =1 cm. C, Seedlings of B73 and *ehsp90.6* at 14 DAG. Scale bar =5 cm. D, Genomic location, structure and mutation site of *ehsp90.6*. *ehsp90.6*, EMS mutant of *hsp90.6.*

*hsp90.6* could be distinguished from the WT as early as 10 days after pollination (DAP), and the plants were light yellow (WT) and light white (*hsp90.6*) (Fig. S2B). The WT and *hsp90.6* grains were separated from the same ear for paraffin embedding. The results showed that the embryo and endosperm development of *hsp90.6* occurred significantly later than those of the WT did, and the subsequent seed filling was significantly hindered in the mutant (Fig. S2E). Therefore, HSP90-6 is involved in the regulation of maize kernel development, as well as seed filling.

### HSP90.6 is a heat shock protein located in the cytoplasm and nucleus

HSP90.6 is a member of the HSP90 subfamily of heat shock proteins and comprises 831 amino acids, with an HATPase_c domain at amino acids 121-283 (Fig. 1D). RT– qPCR analysis showed that *HSP90.6* was expressed in the roots, stems, leaves, tassels and female ears. The transcript level of *HSP90.6* was relatively high in the early stage of grains (Fig. S3B). Then, the expression of *HSP90.6* in kernel subregions was detected by *in situ* hybridization of grains at different stages. *HSP90.6* was highly and evenly distributed in kernels at 5 DAP. The expression in the embryo and endosperm was still strong at 12 DAP, but it was almost undetectable in the seed coat. By 20 DAP, *HSP90.6* transcripts accumulated at low levels except in the scutellum (Fig. 2B). These results showed that *HSP90.6* expression was maintained at a high level in the early stage of grain development. Furthermore, to verify the localization of HSP90.6, subcellular localization was observed, and the results showed that HSP90.6 localized to the nucleus and cytoplasm (Fig. 2C).

**Figure 2.**
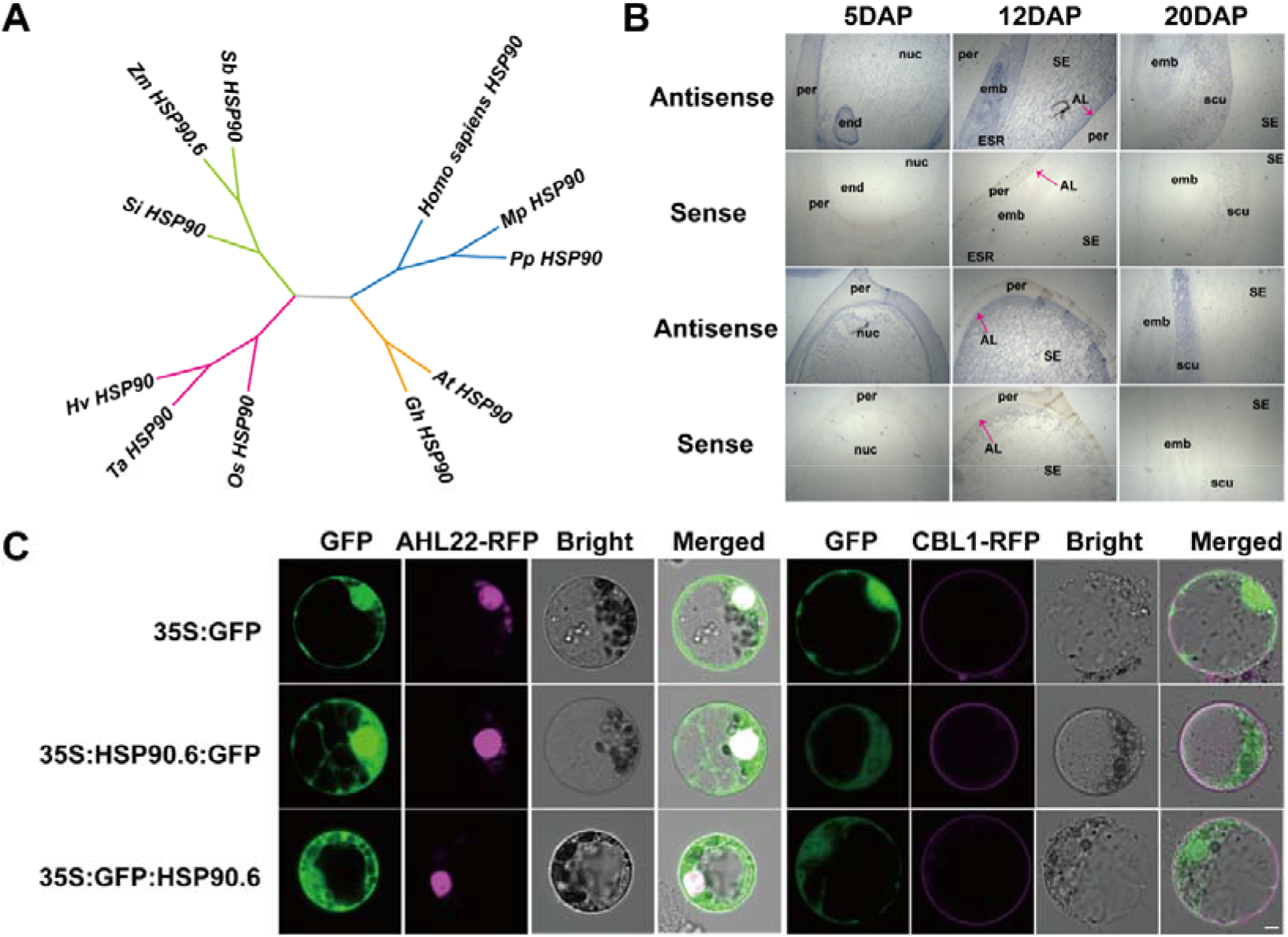
HSP90.6 is a heat shock protein located in the cytoplasm and nucleus A, Phylogenetic relationship of HSP90.6 and its homologs. B, *In situ* hybridization analysis of the expression of *HSP90.6* in LH244 grains at different stages. Scale bar =200 μm. AL, aleurone layer; emb, embryo; end, endosperm; ESR, embryo-surrounding region; nuc, nucellus; per, pericarp; scu, scutellum; SE, starchy endosperm. C, Subcellular localization of HSP90.6 and its colocalization with organelle markers in maize protoplasts. AtAHL22-RFP, nuclear marker; AtCBL1-RFP, cell membrane marker. Scale bar =5 μm.

To understand the evolution of HSP90.6, a phylogenetic tree was constructed based on the HSP90.6 protein sequences of maize and its homologs (Table S2). The results showed that HSP90.6 is especially similar to its orthologs in sorghum and millet (Fig. 2A). Moreover, by analyzing maize HSP family proteins, we found that there are additional small-molecule heat shock proteins (sHSPs). In maize, the HSP90 subfamily has 7 members, including two HSP90.6 members (Fig. S3A), but there are no relevant reports on them.

### A Single-amino acid mutation of HATPase_c affects the function of HSP90.6

The ATPase domain is necessary for HSP90 to function. This domain has ATPase activity and promotes HSP90 to assist protein folding by consuming ATP, which provides energy (Donato and Geisler, 2019; Sima and Richter, 2018). In animals, whether through HSP90 mutation or drug-based inhibition, a reduction in HSP90 activity leads to various abnormal phenotypes (Zuehlke and Johnson, 2010). It has been reported that a single-amino acid mutation in the ATPase domain affects its ATPase activity (Rehn et al., 2020). Therefore, we speculated that the phenotype of smaller grains may be caused by the decreased ATPase activity of *hsp90.6*. To verify this hypothesis, the protein structures of HSP90.6 and *ehsp90.6* were predicted via I-TASSER (https://zhanglab.ccmb.med.umich.edu/I-TASSER/). A structural comparison was conducted by PyMOL (https://pymol.org/), which revealed that the protein structure was not significantly different (Fig. 3A; Fig. S4C). Next, Swiss-Pdb Viewer was used to search for hydrogen bonds between the amino acids, and the results indicated that, unlike Arg127 and Glu123, which formed a hydrogen bond, a hydrogen bond was formed between Gln127 and Val124 of the mutant protein (Fig. S4A and B). In addition, the ligand binding sites and the enzyme active sites were different in *ehsp90.6* (Figs. S5 and S6).

**Figure 3.**
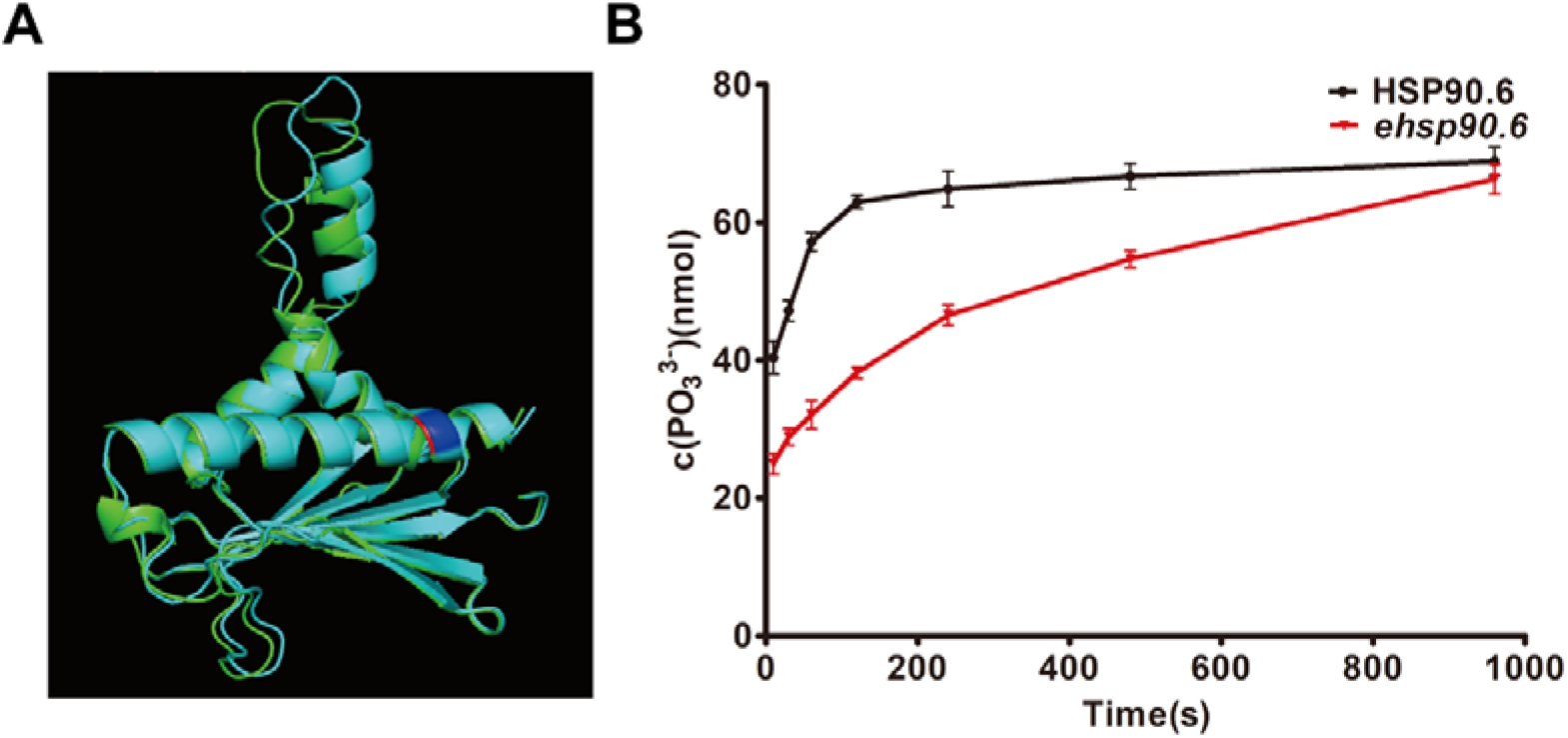
Comparison of HATPase_c structure and verification of its activity in WT and *ehsp90.6*. A, Comparison of predicted WT and *ehsp90.6* HATPase_c domain structures. Green represents HSP90.6, cyan represents *ehsp90.6*, and red and blue represent amino acid mutation sites. B, ATPase activity of HSP90.6 and *ehsp90.6*, determined by the molybdenum blue method. Different reaction times were selected for determination. The error bars for the turnover rates represent ±SDs of three independent measurements (n = 3).

To verify this hypothesis, HSP90.6 and *ehsp90.6* were expressed in *Escherichia coli* to determine ATPase activity. The molybdenum blue method was used to measure the A_660_ after the reaction to obtain the concentration of PO_3_^3-^, reflecting the ATPase activity. The PO_3_^3-^ concentrations at different reaction times were measured and compared, and it was found that, compared with those in the WT, the PO_3_^3-^ concentrations in *ehsp90.6* increased slowly, indicating that the ATPase activity was reduced (Fig. 3B). These results showed that the single-amino acid mutation of *ehsp90.6* reduced its ATPase activity, which may cause the grains of *ehsp90.6* to be smaller than those of the WT.

### The metabolism of carbon and nitrogen is disrupted in *hsp90.6*

The HATPase_c domain is necessary for HSP90, and changes in its activity directly affect the regulatory mechanism in which HSP90 participates (Pearl and Prodromou, 2006). To explore the effect of *hsp90.6* on seed development, WT and *hsp90.6* embryos and endosperm were isolated from ears at 10 DAP, and total RNA was extracted to construct a library for next-generation sequencing. The correlation coefficient of the data was greater than 0.9, indicating that the sequencing quality could be used for further analysis (Fig. S7A and B). Through comparison with the WT data, 6226 differentially expressed genes (DEGs) were identified (screening criteria of |log2(fold-change)|>1 and Q value<0.05), among which 3510 genes were upregulated and 2716 genes were downregulated (Fig. S7C; Table S3).

To understand the cell functions with which the DEGs are involved, Gene Ontology (GO) functional enrichment analysis was conducted, which revealed that HSP90.6 is widely involved in intracellular processes, including protein homeostasis, signal transduction, transcriptional regulation and nutrient metabolism. As expected, genes related to protein metabolism were highly enriched (Fig.4A; Table S4), because HSP90 mediates protein folding or assembly and helps polypeptides fold and be transported (Schopf et al., 2017). Regarding signal transduction and transcriptional regulation, related studies have reported that HSP90 carries out these functions by activating steroid hormone receptors, protein kinases and transcription factors (Picard et al., 1990; Pratt, 1997; Jackson et al., 2004).

**Figure 4.**
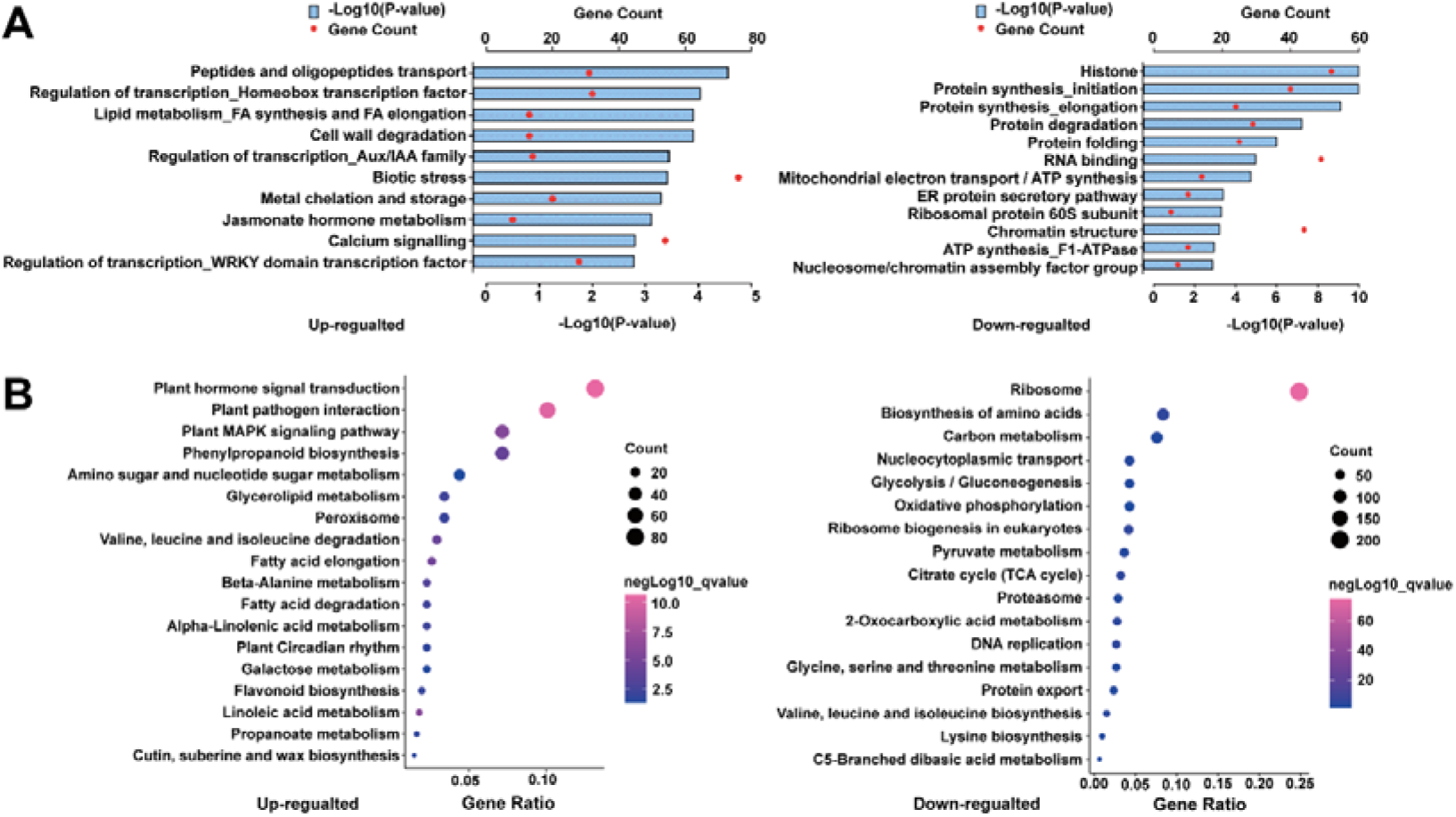
Effects of *hsp90.6* on kernel development A, Functional classification of DEGs between *hsp90.6* and WT. B, KEGG pathway enrichment analysis of the DEGs.

For nutrient metabolism, Kyoto Encyclopedia of Genes and Genomes (KEGG) pathway analysis was performed on the DEGs to determine the metabolic processes in which they participated. Pathways involving biosynthesis of amino acids and carbon metabolism were highly enriched in downregulated genes (Fig. 4B; Table S5). This means that the grain filling of *hsp90.6* was severely affected. Therefore, the DEGs affecting grain filling were analyzed. The transcript levels of *O2* (*Opaque 2*), *O11* (*Opaque 11*), *OS1* (*Opaque Endosperm and Small Germ 1*), *MN6* (*Miniature Seed 6*), *Bt2* (*Brittle 2*), *PBF1* (*Prolamin-box factor 1*), *NKD2* (*Naked endosperm 2*), and *NAC130* (*NAM, ATAF, and CUC TFs 130*) decreased significantly. O11 is an endosperm-specific transcription factor that can transcriptionally activate *NKD2*, which directly regulates the expression of *O2*. O2 is the central regulator of grain filling, which, through interaction with PBF1, can transcriptionally activate sucrose cleavage, starch biosynthesis, and protein storage-related genes (Deng et al., 2020; Feng et al., 2018; Gontarek et al., 2016; Song et al., 2019; Yang et al., 2016; Yang et al., 2021; Yi et al., 2021; Zhang et al., 2015, 2016). NAC130 regulates the expression of *Bt2* and 16-kDa γ-zein. The starch and protein contents of *nac130* seeds are dramatically reduced (Zhang et al., 2019).

### HSP90.6 affects nutrient metabolism by interacting with RPN6, PBD2 (PRC2) and GF14-6

To explore how HSP90.6 affects carbon and nitrogen metabolism, an antibody against HSP90.6 was constructed (Fig. S8), and total proteins of WT and *hsp90.6* embryos and endosperm at 10 DAP were extracted for immunoprecipitation (IP). The immunoprecipitated proteins were identified by mass spectrometry (MS), after which their sequences were queried via BLAST in the MaizeGDB. After comparing the IP-MS results of the WT and *hsp90.6*, we obtained a total of 52 specific proteins and the corresponding gene descriptions and studies were checked in Ensembl (http://ensembl.gramene.org/Zea.mays) (Fig. 5A and B; Table S6).

**Figure 5.**
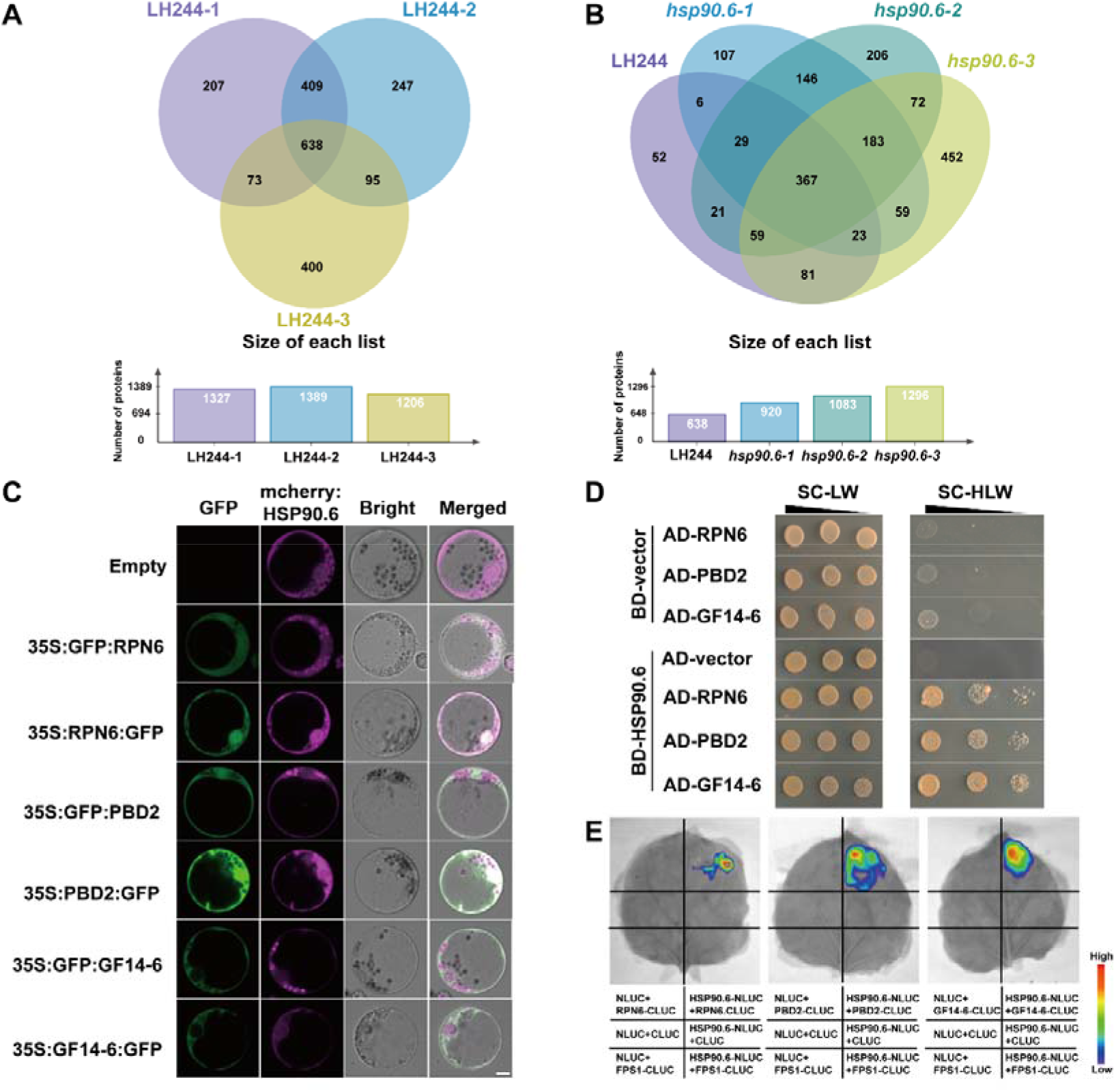
Screening, colocalized and interaction of HSP90.6-interacting proteins A, Venn diagram of IP-MS results of three biological repeats of HSP90.6. B, Venn diagram of IP-MS results of three biological repeats of HSP90.6 overlapping with three biological repeats of *hsp90.6*. C, Subcellular localization showing that HSP90.6 colocalize with PRN6 and PBD2 and overlaps with GF14-6. Scale bar =5 μm. D, Yeast two-hybrid assays showing that HSP90.6 interacts with PRN6, PBD2 and GF14-6. AD, activating domain; BD, binding domain. E, LCI showing that HSP90.6 interacts with PRN6, PBD2 and GF14-6. The fluorescent signal intensities represent their interaction activities. FPS1 (Farnesyl Pyrophosphate Synthetase 1), negative control.

Considering the connection between nitrogen recycling and the ubiquitin– proteasome system, the 26S proteasome subunits RPN6 and PBD2 (PRC2) were selected for verification. The interaction between HSPs and the 26S proteasome has been reported in animals and yeasts (Kandasamy and Andréasson, 2018; Park et al., 2007; Verma et al., 2000). However, there are no reports of embryonic lethality caused by the loss of function of the 26S proteasome (Wang et al., 2019). GF14-6, another protein identified via IP-MS, attracted our attention. The GF14-6 protein mainly regulates C/N metabolism (Diaz et al., 2011), and according to the KEGG results, genes involved in carbon metabolism-related were downregulated (Fig. 4B). To verify the relationships between HSP90.6 and these proteins, the subcellular localization of RPN6, PBD2 (PRC2) and GF14-6 was predicted through CELLO (http://cello.life.nctu.edu.cn). RPN6 and PBD2 (PRC2) were located in the nucleus and cytoplasm, which is consistent with the location of HSP90.6. GF14-6 was mainly distributed in the cytoplasm, which overlapped with the location of HSP90.6. Subsequently, subcellular localization was performed in maize protoplasts, and RPN6 and PBD2 (PRC2) completely overlapped with HSP90.6. The cytoplasmic location of GF14-6 also overlapped with that of HSP90.6 (Fig. 5C). In summary, HSP90.6 colocalizes with RPN6, PBD2 (PRC2) and GF14-6. In addition, yeast two-hybrid and luciferase complementation imaging (LCI) assays were conducted to prove the interactions between HSP90.6 and RPN6, PBD2 (PRC2) and GF14-6 (Fig. 5D and E). These results revealed that HSP90.6 interacted with RPN6, PBD2 (PRC2) and GF14-6 both *in vivo* and *in vitro*.

### Loss of function of HSP90.6 reduces 26S proteasome activity and affects nitrogen recycling

In eukaryotes, the autophagy–lysosome system and ubiquitin–proteasome system (UPS) are responsible for nitrogen recycling by the degradation of short-lived regulatory proteins and soluble misfolded proteins, and the core component responsible for this activity is the 26S proteasome (Finley et al., 2012; Marshall and Vierstra, 2019; Samant et al., 2018; Schubert et al., 2000; Vierstra, 2009). In addition to proteasome degradation requiring the complex structure of the complete holoenzyme, specific ubiquitin labeling of suitable degradation substrates is needed (Budenholzer et al., 2017; Bard et al., 2018). HSP90 has a large substrate-binding interface, which preferentially binds to late-folding intermediates that have containing higher order structures. HSP90 activity is usually part of the final step to promote protein folding, assembly of multiprotein complexes, and binding of ligands to substrates (Abildgaard et al., 2020; Jakob et al., 1995; Karagöz et al., 2014). As a subunit of RP, RPN6 is an important component of the 26S proteasome and has been identified as a molecular tweezer that fixes CP and RP together to form a fully activated 26S proteasome (Cho et al., 2015; Estrin et al., 2013; Isono et al., 2005; Pathare et al., 2012; Santamaria et al., 2003; Tian and Trader, 2020). Specifically, RPN6 interacts with RPT6 of RP and the α2 subunit of CP (Chen et al., 2016; Tian and Trader, 2020). PBD2 (PRC2) is a subunit of CP and is responsible for endowing active sites with chymotrypsin cleavage properties and maintaining the degradation activity of the 26S proteasome (Arendt and Hochstrasser, 1997; Dick et al., 1998; Heinemeyer et al., 1997; Kisselev et al., 1999, 2003; Kumar Deshmukh et al., 2019; Marshall and Vierstra, 2019). We speculated that the interactions of HSP90.6 with RPN6 and PBD2 (PRC2) would affect the function of the 26S proteasome.

To verify that the function of the 26S proteasome was impaired, the 26S proteasomes were separated via molecular sieves from WT and *hsp90.6* embryos and endosperm at 18 DAP. A total of 20 fractions were collected and detected by Western blotting with anti-HSP90.6/PSMD13/PSMA3. The results showed that 26S proteasomes were distributed in fractions 2-7 (Fig. 6A). Then, fractions 2-7 were mixed, and a specific fluorescent matrix was used to detect the hydrolytic activity of the 26S proteasome in WT and *hsp90.6* on Suc-LLVY-AMC. Compared with that of the WT, the hydrolysis activity of the 26S proteasome of *hsp90.6* was reduced by 74.9% (Fig. 6B). At the same time, native PAGE was used to test the activity of the 26S proteasome, and consistent results were obtained (Fig. 6C). Therefore, *hsp90.6* presented reduced 26S proteasome activity.

**Figure 6.**
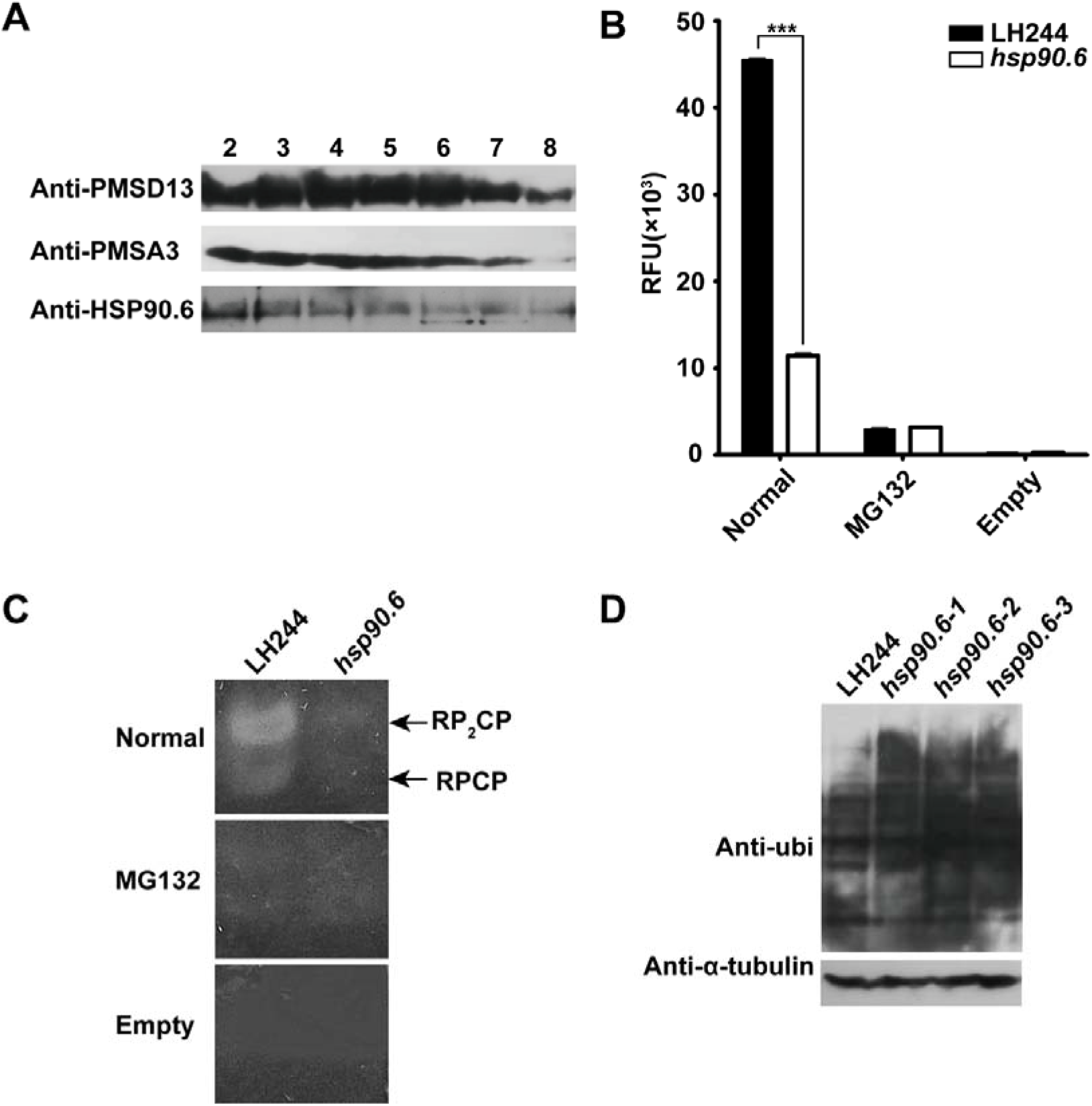
*hsp90.6* reduces 26S proteasome activity A, Total protein extracted from the embryo and endosperm at 18 DAP were separated by a molecular sieve and then analyzed by Western blotting in conjunction with anti-HSP90.6, anti-PSMD13 and anti-PMSA3. PSMD13 encodes a non-ATPase subunit of the 19S regulator. PMSA3 encodes a 20S core α subunit. B, 26S proteasome activity analysis of embryos and endosperm at 18 DAP in solution. RFU, relative fluorescence units; Normal, 26S proteasome and Suc-LLVY-AMC; MG132, 26S proteasome, Suc-LLVY-AMC and MG132; Empty, Suc-LLVY-AMC. For each sample, three independent biological replicates were analyzed. Error bars = average values ± SDs. n = 3. ***, P<0.001 (Student’s t test); C, 26S proteasome activity in-gel analysis of embryos and endosperm at 18 DAP; D, Ubiquitinated proteins of embryos and endosperm at 18 DAP were analyzed by Western blotting in conjunction with anti-ubi.

As proteasome damage increases the level of ubiquitinated substrate, anti-ubi was used to analyze the level of ubiquitinated proteins in the embryo and endosperm of WT and *hsp90.6*. The Western blot results showed that the amount of ubiquitinated proteins of *hsp90.6* obviously increased (Fig. 6D). Next, IP-MS was used to analyze ubiquitinated proteins in the embryo and endosperm at 18 DAP. Through the comparison of WT and *hsp90.6*, a total of 271 accurately quantified proteins were identified. Under the conditions of |log2(fold-change)|≥1 and Q value<0.05, 26 differentially expressed proteins were screened. The accumulation of 20 proteins was increased, which included mainly 60S ribosomal proteins and enzymes. 60S ribosomal proteins are related to translation, and the enzymes are involved in different processes. For example, Zm00001d011139 (*Histone Deacetylase 102*) regulates histone acetylation, and its accumulation disrupts gene expression (Shahbazian and Grunstein, 2007). The increase in proteins further indicated that the activity of the 26S proteasome decreased. However, the expression of 6 proteins was downregulated, which may be because of the failure of the transcriptional regulation pathway involved in HSP90.6 (Table S7). The results suggested that deletion of HSP90.6 reduced 26S proteasome activity, resulting in the accumulation of ubiquitinated proteins; these accumulated proteins cannot be recycled, thus affecting nitrogen recycling. This is consistent with the downregulation of amino acid synthesis-related genes according to the KEGG analysis.

### HSP90.6 interacts with GF14-6 to influence the carbon metabolism

Among the DEGs, those related to carbon metabolism were significantly downregulated in *hsp90.6*, which could have an impact on grain filling. Although the genes encoding the core transcription factors *O2*, *O11*, *NKD2*, and *NAC130* that regulate grain filling were all downregulated in *hsp90.6* (Table S3), the absence of these proteins did not completely hinder grain filling, indicating that there were other proteins involved in regulating of regulating grain filling. GF14-6 is a subtype of 14-3-3 proteins and is ubiquitous in all eukaryotic cells. 14-3-3 proteins are typical cytoplasmic proteins and participate in various pathways such as sugar metabolism and lipid metabolism (Aducci et al., 2002). It has been reported that 14-3-3 proteins play a direct role in the process of starch accumulation (Sehnke et al., 2002). In addition, there is evidence in maize that *ZmGF14-4* and *ZmGF14-6* are involved in glycolysis and post glycolysis metabolism (Dou et al., 2015). 14-3-3 proteins usually combine with phosphorylated proteins to function, and PTM Viewer (https://www.psb.ugent.be/webtools/ptm-viewer) showed that both Ser677 and Ser688 of HSP90.6 are phosphorylated (Fig. S9).

To understand the regulatory mechanism of GF14-6, IP-MS was performed. A 35S:GF14-6:GFP vector was constructed and transformed into maize protoplasts. After 24 h of culture, the protoplasts were collected and the total proteins were extracted. Anti-GFP was used for IP-MS, and 25 candidate proteins were obtained (Fig. 7A and B). Next, we predicted the subcellular localization of these 25 proteins in CELLO and finally ultimately identified 9 proteins localized in the cytoplasm (Table S8). We selected the genes related to glucose metabolism, including Zm00001d015376 (*Phosphoglycerate kinase* (*GPC1*)), Zm00001d046170 (*Phosphoenolpyruvate carboxylase 1* (*PEP1*)) and Zm00001d049641 (*Glyceraldehyde-3-phosphate dehydrogenase* (*PGK3*)) for verification. Moreover, we found that Zm00001d049239 (*Adenosyl homocysteine hydrolase1* (*AHH1*)) was the only protein whose localization overlapped with that of HSP90.6 according to the IP-MS results, which can catalyze the hydrolysis of SAH and enable smooth methyl transfer (Min et al., 2014). The interaction between GF14-6 and these 4 proteins was verified with LCI. The obtained results preliminarily confirmed that GF14-6 interacts with GPC1, PGK3 and AHH1 (Fig. 7C), which implied that HSP90.6 interacts with GF14-6 to participate in grain filling via carbon metabolism.

**Figure 7.**
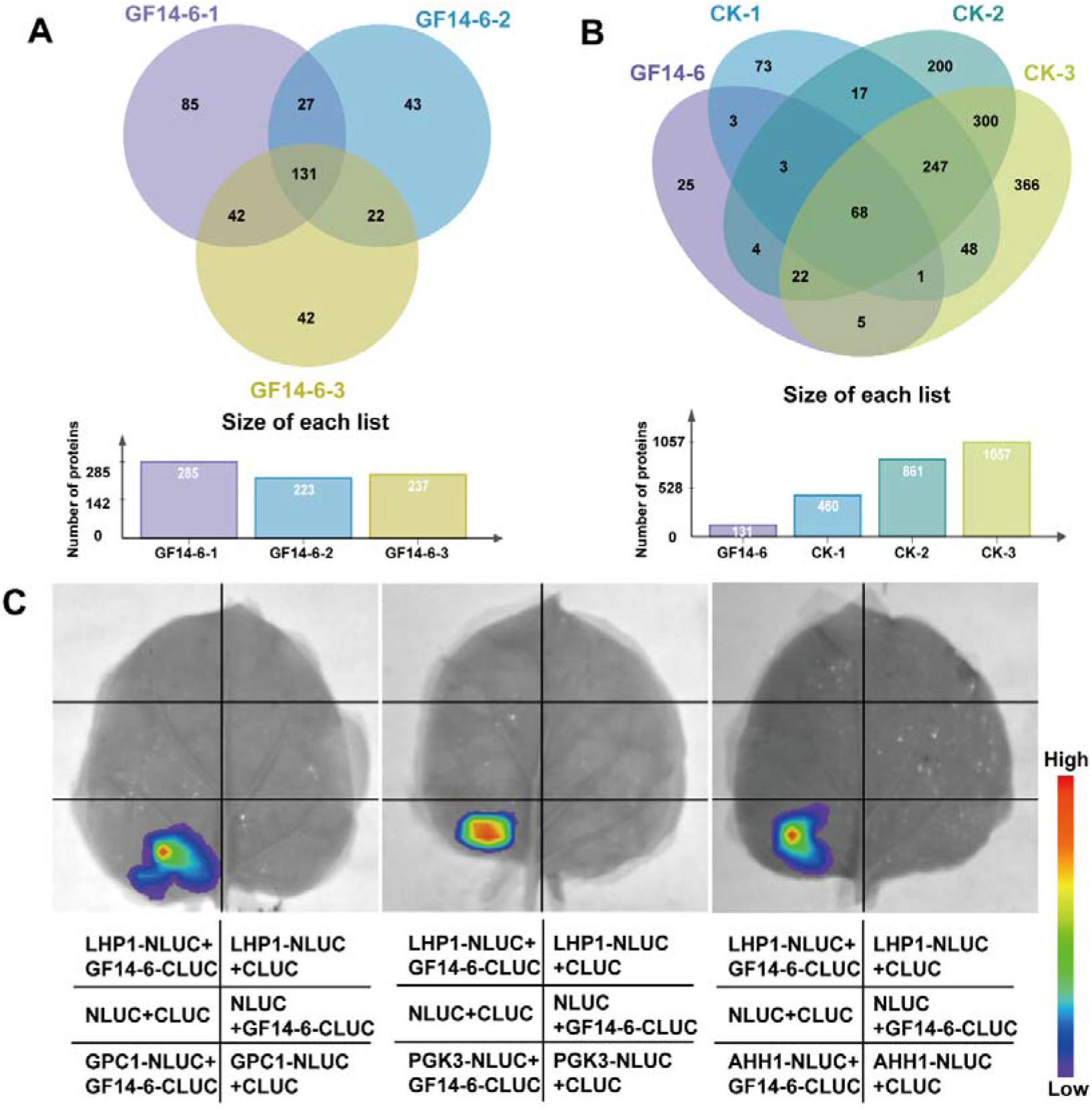
Identification and verification of GF14-6 interaction proteins. A, Venn diagram of the IP-MS results of three biological repeats of GF14-6. B, Venn diagram of the IP-MS results of three biological repeats of GF14-6 overlapping with three biological repeats of the control. GF14-6 was expressed in protoplasts via 35SS:GF14-6:GFP; control protoplasts were transformed with 35S:GFP. C, LCI showing that GF14-6 interacted with GPC1, PGK3 and AHH1. LHP1 (Like Heterochromatin Protein 1), negative control.

## Discussion

Carbon and nitrogen are essential nutrients for plant growth, and their metabolism in plants determines the morphogenesis of plant organs (Goel et al., 2016). In this work, we found that HSP90.6 was involved in carbon and nitrogen metabolism, suggesting that it functions in maintaining nutrient metabolism in maize kernels. *hsp90.6* showed severe kernel development defects (Fig. S2B-D), which indicated that the HSP90.6 protein participates in the regulation of maize kernel development. There are reports that the loss of AtHSP90.6 function in *Arabidopsis* can cause embryonic death, and the mutation of chloroplast-targeted AtHSP90.5 resulted in embryonic death (Feng et al., 2013; Luo et al., 2019). However, the underlying molecular mechanism remains unknown. HSP90.6 interacted with RPN6/PBD2 (PRC2) to regulate nitrogen recycling in the cell by influencing 26S proteasome activity. It has been reported that AtHSP101 works synergistically with the 26S proteasome to promote the clearance of ubiquitinated proteins by refolding rather than proteasomal degradation, and AtHSP101 deletion has no effect on 26S proteasome (McLoughlin et al., 2019). Therefore, the regulatory mechanisms of HSP90.6 and AtHSP101 are different. HSP90.6 interacted with GF14-6 to regulate carbon metabolism and affect grain filling (Fig. 8).

**Figure 8.**
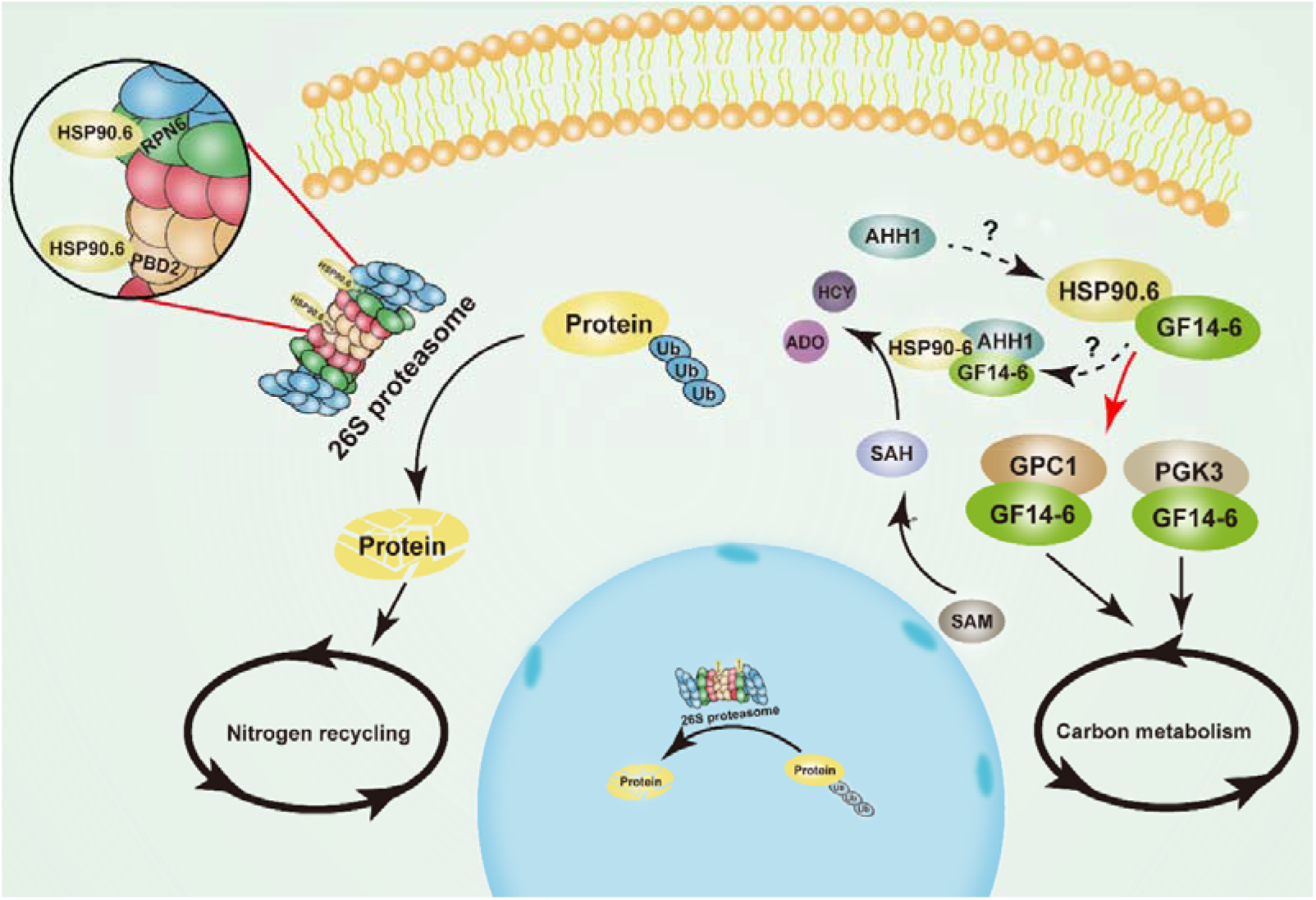
Model of ZmHSP90.6 regulating nutrient metabolism in maize kernel development. HSP90.6 interacts with RPN6/PBD2 (PRC2) to regulate 26S proteasome activity and nitrogen recycling. In addition, HSP90.6 interacts with GF14-6 to regulate PGK3 and GPC1 to participate in carbon metabolism. HSP90.6 may form a trimer with GF14-6 and AHH1 to participate in SAH hydrolysis. The red arrow indicates the molecular mechanism revealed in the present work.

### HSP90.6 is required for 26S proteasome-mediated nitrogen recycling

Nitrogen recycling is inseparable from the ubiquitin proteasome system, and the core of the ubiquitin proteasome system is the 26S proteasome. The assembly of the 26S proteasome requires a series of proteasome-specific chaperones to ensure that each subunit finds its correct position in the newly synthesized proteasome (Budenholzer et al., 2017). Many related chaperones have been reported in animals, such as PAC1-PAC2 and PAC3-PAC4 heterodimers to promote the assembly of α loops (Kusmierczyk et al., 2008, 2011; Yashiroda et al., 2008); Ump1 plays a role in the binding of β subunits and half-dimers (Ramos et al., 1998). Here, we proved that the loss of HSP90.6 results in decreased 26S proteasome activity and the accumulation of ubiquitinated proteins that affect nitrogen recycling (Fig. 6B and C; Table S7). HSP90 has a large substrate-binding interface, which preferentially binds late-folding intermediates containing higher order structures (Amanda et al., 2020; Jakob et al., 1995; Karagöz et al., 2014). RPN6 has been identified as a molecular tweezer that fixes CP and RP together to form a fully activated 26S proteasome. We speculated that HSP90.6 may promote the function of RPN6 as a molecular tweezer. PBD2 (PRC2) is the core component of 26S proteasome function (Arendt and Hochstrasser, 1997; Dick et al., 1998). The loss of HSP90.6 may cause PBD2 (PRC2) to lose its chymotrypsin cleavage activity, which directly affects the function of the 26S proteasome.

### HSP90.6 plays an important role in carbon metabolism

In a study on *ZmDEK40*, the lack of DEK40 significantly affected the biogenesis of 20S CP, resulting in a decrease in 26S proteasome activity (Wang et al., 2019). However, *dek40* produced smaller kernels only, suggesting that the interaction of HSP90.6 with RPN6 and PBD2 (PRC2) was insufficient to lead to the lethal phenotype of *hsp90.6*. RNA sequencing (RNA-seq) analysis showed that the loss of HSP90.6 caused the downregulation of carbon metabolism-related genes (Fig. 4B), and the 14-3-3 protein GF14-6 was identified by IP-MS (Table S6).

In plants, 14-3-3 proteins regulate protein interactions and enzyme activity in a phosphorylation-dependent manner, those involving transcription factors, ion channel proteins, and kinases (Kong and Ma, 2018). Phosphorylation of HSP90 has been studied in animals. A specific phosphorylation site of human HSP90β, Ser365, has been shown to affect the interaction of HSP90 and CDC37 (Nguyen et al., 2017). In addition, a large number of phosphorylation sites on HSP90α and HSP90β have been predicted by the PhosphositePlus database (Hornbeck et al., 2012), which also implied the possibility of interactions between HSP90 and 14-3-3s. Proteomic analysis has revealed that 14-3-3s are involved in glycolysis, fatty acid synthesis, protein storage and other C/N metabolic pathways (Dou et al., 2015; Zhang et al., 2019). We found that the 14-3-3 protein GF14-6 interacted with the carbon metabolism-related proteins GPC1 and PGK3 (Fig. 7A-C). Therefore, we speculated that the loss of function of HSP90.6 led to impaired carbon metabolism and severely inhibited in grain filling. However, this was indirect, and HSP90.6 may be phosphorylated first, and then combined with GF14-6. Their combination may cause conformation changes in GF14-6, which would become active and start to activate GPC1 and PGK3. Notably, through IP-MS, we found that HSP90.6 and GF14-6 had an overlapping protein (AHH1), which is responsible for catalyzing the decomposition of adenosine homocysteine to ensure the normal supply of methyl compounds (nucleic acid, protein). Therefore, we speculated that the process may be carried out by the formation of a complex comprising HSP90.6, GF14-6 and AHH1.

The genes encoding the transcription factors *O2*, *O11*, *NKD2*, and *NAC130*, which have been reported to regulate grain filling, were all downregulated in *hsp90.6*, which meant that HSP90.6 may play an upstream regulatory role. However, HSP90.6 had no transcriptional activation activity (Fig. S10), so this regulation occurred indirectly. As its regulatory process is still unknown, an abundance of research is still needed.

## Materials and Methods

### Plant material

Mutations in kernels were induced by EMS. For this, mature pollen grains of inbred line B73 were soaked in 0.015% EMS solution (volumetric ratio of 1:10) for 10-15 min, and then selfed to obtain T1 seeds. The T1 seeds were planted and selfed to obtain the T2 generation for phenotypic identification.

The *hsp90.6* knockout mutant was obtained by clustered, regularly interspaced, short palindromic repeats (CRISPR)/CRISPR-associated 9 (Cas9) and transformed by the maize genetic transformation platform of the National Maize Improvement Center of China Agricultural University. The approximate operation method was as follows: two gRNA sequences were inserted into the binary vector pXUE411C-BG according to a previous report (Xing et al., 2014), and gRNAs from CRISPR-P (http://crispr.hzau.edu.cn/CRISPR/), CCCGGAGCAGCTCATTCCTA and TCAATGAGTACTTGGGCCCT, were selected. Prepared *Agrobacterium* cells containing the target vector were infiltrated into LH244 embryos (1.5-1.8 mm) at 12 DAP for *Agrobacterium*-mediated transformation (Lagrimini, 2018). Herbicides were used select positive transformants, which were verified by DNA sequencing at the seedling stage. In this work, the embryos and endosperm for RNA-seq, IP-MS and 26S proteasome experiments were purified using CRISPR/Cas9 materials.

### BSR

T2 seeds of WT (B73) and EMS mutant embryos and endosperms at 12 DAP were taken and analyzed by BSR. Total RNA was extracted with TRIzol^TM^ reagent (15596018, Thermo Fisher Scientific, China).

cDNA libraries were constructed using an AHTS Stranded mRNA-seq Library Prep Kit for Illumina V2 (NR612, Vazyme, China). The sequences were obtained by an Illumina X-10 instrument (Illumina, America). The raw data were compared with the B73_V4 reference genome data by HISAT2 v2.0.4 (http://daehwankimlab.github.io/hisat2/), with the default parameters (HISAT is a fast splice aligner with low memory requirements); the BAM file was output; and SAMtools v0.1.19 (http://samtools.sourceforge.net/) (in the sequence alignment/map format) was used for sorting. Single-nucleotide polymorphism (SNP) calling was performed with unique reads through SAMtools v0.1.19 and BCFtools. The screening criteria for high-confidence SNPs were as follows: 1) SNPs supported at least five reads, and 2) SNPs in the WT pool corresponding to the mutant pool were detected, and use SnpEff v4.2 (https://pcingola.github.io/SnpEff/), which is a program for annotating and predicting the effects of SNPs, was used to annotate the SNPs.

### Identification of candidate genes

The degree of linkage between the frequency of each SNP and the phenotype was calculated, and the SNP-corresponding genes with the highest linkage degree were screened. Then, SNPs in on the genes of 300 segregating populations of the T3 generation were detected, and candidate genes that cosegregated with the phenotype were identified.

### qRT–PCR

Five micrograms of Total RNA was collected, and a Hifair® II 1st Strand cDNA Synthesis Kit (with gDNA Digester Plus) (11121ES60, YEASEN, China) was used for reverse transcription. qRT–PCR was performed in conjunction with Takara TB Green® Premix Ex Taq™ (Tli RNase-H Plus), Bulk (RR420L, Takara, Japan).

### Phylogenetic analysis

The protein sequence of HSP90.6 was searched by BLASTP in NCBI database (https://www.ncbi.nlm.nih.gov/) and a phylogenetic tree was constructed with MEGA v7.0 (www.megasoftware.net). The Muscle algorithm was used for multiple sequence alignment. The statistical method used was the adjacency matrix method (neighbor-joining (NJ) method). The bootstrap method was used for the phylogenetic tree tests, in which the number of tests was 1000 and the replacement model was the Poisson model. The phylogenetic tree was visualized by iTOL (https://itol.embl.de/).

### Subcellular localization

The coding sequence (CDS) of *HSP90.6* was cloned into the N-terminal region and C-terminal region of pUC:35S:GFP, and the resulting expression vectors were transformed into maize protoplasts using PEG-Ca^2+^ as described by Yoo et al (Yoo et al., 2007). Then, the protoplasts were cultivated at 28°C in the dark for 18-24 h. Images were taken with a confocal microscope (Zeiss 710, Germany) that showed the fluorescence signal of GFP under excitation at 488 nm and that of RFP under excitation at 561 nm.

### ATPase activity

The CDSs of the *HSP90.6* and *ehsp90.6* were ligated into pET-32a, which were subsequently transformed into BL21 (DE3) competent cells for protein expression and purification. ATPase activity was measured by the molybdenum blue method with minor modifications (Rodriguez et al., 1994; Yamane et al., 2019). We incubated 75 μL of reaction solution (30 mM Tris at pH 7.5, 8 mM MgCl_2_, 100 mM NaCl, 1 mM dithiothreitol (DTT), 4% (w/v) sucrose, 30 μg/ mL bovine serum albumin (BSA), 1 mM phenylmethylsulfonyl fluoride (PMSF), 1.5 μL of 100 nM ATP and 100 nM of prokaryotic purified HSP90.6/*hsp90.6* at 30°C for 10 s/30 s/1 min/ 2 min/4 min/8 min/16 min and then immediately added 175 μL of color developing solution (0.35% (m/v) (NH_4_)_2_MoO_4_.4H_2_O, 0.86 mol/L H_2_SO_4_, 1.4% (w/v) ascorbic acid). Then, they were incubated at 42°C for 30 min, and the absorbance (A_660_) was measured with a micro plate reader (Molecular Devices i3x, America).

### RNA-seq analysis

A cDNA library was constructed with a Hieff NGS MaxUpTM Dual-mode mRNA Library Prep Kit for Illumina (H9001360, Vazyme, China) and sequenced on an Illumina Nova-seq platform (Illumina, America), generating 150-nucleotide paired-end reads. FASTP v0.20.1 (http://github.com/OpenGene/fastp) was used to remove low-quality reads. HISAT2 v2.2.1 (http://daehwankimlab.github.io/hisat2/) with the default parameters were used to map the original reads with the sequence information of the B73_V4 reference genome. Then the gene expression levels were measured via fragments per kilobase of exon model per million mapped fragments (FPKM) using Cuffflinks (V2.2.1). The differential gene screening criteria were |log2(fold-change) |>=1 and Q value <0.05.

### IP assays

Polyclonal anti-HSP90.6 antibodies were produced in white rabbits by HuaBio (China, http://www.huabio.com). The peptide sequence used as an antigen was ASIPPRPASNGAPGC.

The embryos and endosperm of WT (LH244) and *hsp90.6* at 12 DAP were taken, and total protein was extracted according to the manufacturer’s instructions of a plant protein extraction kit (KGP750, KeyGEN BioTECH, China). The reagents used in the IP were from a BEAVER BeaverBeads^TM^ Protein A (or A/G) Immunoprecipitation Kit (22202-20, Beaverbio Medical Engineering, China). The proteins obtained by IP were detected by Western blotting and identified by MS. The MS identification was performed by staff at the Biomass Spectrometry Laboratory of the School of Biology, China Agricultural University.

### Yeast two-hybrid assays

The CDS of *HSP90.6* was linked downstream of the GAL4 BD domain of pGBKT7. RPN6, PBD2 and GF14-6 were inserted into pGADT7. An EX-Yeast Transformation Kit and related media were used (ZC135, ZOMANBIO, China).

### LCI assays

The CDSs of *HSP90.6*, RPN6, PBD2, and GF14-6 were inserted in pCAMBIA1300-nLUC and pCAMBIA1300-cLUC. Then, the resulting plasmids were transformed into *Agrobacterium* EHA105 competent cells, and the transformants were injected into leaves of 3-week-old *Nicotiana benthamiana* seedlings. After incubation at 28°C for 48-72 h, the leaves were removed, and a 1 mM solution of D luciferin (40902ES01, YEASEN, China) was sprayed onto the leaves. The fluorescence signal was captured by a NightShade imaging system (Berthold LB985, Germany).

### In-gel peptidase activity assays of the 26S proteasome

Embryos and endosperm of WT (LH244) and *hsp90.6* were at 18 DAP used for 26S proteasome purification (Book et al., 2010; Li et al., 2015; Wang et al., 2019).

Total protein was extracted and subjected to fast-performance liquid chromatography (FPLC) using a Superdex Increase 200 10/300GL column (10246670, GE Healthcare, Sweden) to collect 1 mL fractions per sample. Twenty samples from the fraction collector were placed on ice as soon as the run was completed to detect the proteasome complexes by Western blotting with anti-HSP90.6, anti-PSMD13 and anti-PSMA3 (A6956 and A1245, ABclonal, America).

The proteasome complexes (2-7, ≥670 kD) of the WT and *hsp90.6* at 18 DAP were mixed, and 0.5 μL (100 nm) was taken. Afterward, 9.5 μL of buffer A (50 mM Tris-HCl at pH 7.5, 10% (v/v) glycerol, 5 mM ATP, 5 mM MgCl_2_, 1 mM DTT and 1 mg/mL creatine phosphokinase) and 2.5 μL of 5X native gel loading buffer were added. A 4% native gel that included 1 mM ATP was used to separate 26S proteasome proteins at 4°C and 100 V for 3.5 h. Next, the gel was rinsed with water, after which 10 mL of development buffer (50 mM Tris at pH 7.5, 150 mM NaCl, 5 mM MgCl_2_, 1 mM ATP and 100 μM Suc-LLVY-AMC) was added, followed by incubation at 30°C at 30 rpm for 30 min. The gel was subsequently scanned at 365 nm by an UV irradiator (Bio-Red, America)

### In-solution peptidase activity assay for 26S proteasomes

Buffer A was replaced with 100 μL of development buffer, and 35 pmol of 26S proteasome proteins were added, followed by incubation at 30°C for 30 minutes. One milliliter of 1% SDS was used to stop the reaction. The fluorescence was measured at 380 nm excitation light and 460 nm emission light by a multifunctional microplate reader (Tecan Spark, Switzerland).

### Ubiquitylome analysis

The ubiquitylome experiment was carried out for two independent biological replicates, and each replicate consisted of a mixture of embryo and endosperm from different transformants at 18 DAP. A Ubiquitin Detection Kit (SMQ-SKT-131-20, StressMarq, Canada) was used for IP. The ubiquitinated proteins were analyzed by MS and MaxQuant v1.3.0.5 (https://www.maxquant.org/) and compared with the sequence information housed in the MaizeGDB database. Proteins were considered differentially expressed under |log2(fold-change) |>=1 and Q value <0.05.

### Accession numbers

HSP90.6 (Zm0000d041719); O2 (Zm00001d018971); O11 (Zm00001d003677); OS1 (Zm00001d002661); MN6 (Zm00001d037926); Bt2 (Zm00001d050032); PBF1 (Zm00001d005100); NKD2 (Zm00001d026113); NAC130 (Zm00001d008403); RPN6 (Zm00001d041456); PBD2 (Zm00001d013505); GF14-6 (Zm00001d031688); FPS1 (Zm00001d009431); LHP1 (Zm0000d014449); AtAHL22 (At2g45430); AtCBL1 (At4g17615).

## Supplemental Data

**Supplemental Figure S1.** Phenotypic statistics of *ehsp90.6*.

**Supplemental Figure S2.** Phenotypic tests of the ears of the CRISPR/Cas9-generated mutant *hsp90.6*.

**Supplemental Figure S3.** Characteristics of *HSP90.6*.

**Supplemental Figure S4.** Prediction and comparison of protein structure

**Supplemental Figure S5.** Structural prediction of HSP90.6.

**Supplemental Figure S6.** Structural prediction of *ehsp90.6*.

**Supplemental Figure S7.** Analysis of DEGs.

**Supplemental Figure S8.** Western blot analysis of the specificity of anti-HSP90.6.

**Supplemental Figure S9.** PTM Viewer results of HSP90.6.

**Supplemental Figure S10.** Analysis of transcriptional activation activity of HSP90.6

**Supplemental Table S1.** Candidate genes identified in the EMS mutants.

**Supplemental Table S2.** Phylogenetic tree related gene information.

**Supplemental Table S3.** Genes differentially expressed between *hsp90.6* and the WT.

**Supplemental Table S4.** Functional classification of DEGs between *hsp90.6* and the WT.

**Supplemental Table S5.** KEGG pathway enrichment analysis of the DEGs.

**Supplemental Table S6.** IP-MS data of HSP90.6.

**Supplemental Table S7.** Quantitative analysis of ubiquitinated proteins.

**Supplemental Table S8.** IP-MS data of GF14-6.

**Supplemental Table S9.** Primers used in this research.

## Acknowledgments

This work was supported by the National Natural Science Foundation of China (91935303; 31971957; 31971959; 91935305), and National Key R&D Program of China (2021YFD1200700).

## Author contributions

J.X., Z.Y. and M.Z. designed the experiments. J.X., Z.Y. and X.F. performed the experiments. J.X., W.S., Q.Z. and J.L. wrote the manuscript. Y.C. and X.Z. screened the mutant *hsp90.6.* K.T. analyzed the RNA-seq data. L.E. and H.Z. revised the manuscript. The authors declare that there are no conflicts of interest.

## Data availability

All the data supporting the findings of this research are available within the paper.

---

**Figure S1.**
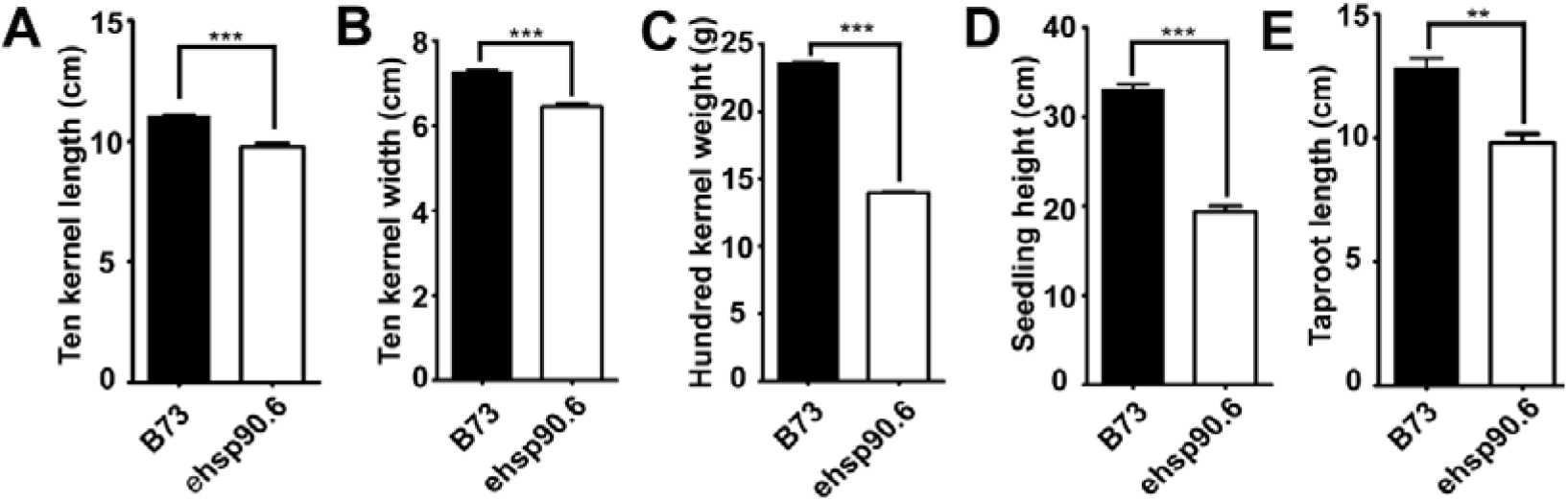
Phenotypic statistics of *ehsp90.6* A, Length of ten seeds of B73 and *ehsp90.6*. B, Width of ten seeds of B73 and *ehsp90.6*. C, Weight of one hundred seeds of B73 and *ehsp90.6*. Three independent biological replicates were included for each sample. n=3 replicates. Error bars = 3 average values ± SDs. ***, P<0.001 (Student’s t test). D, Average height of B73 and *ehsp90.6* plants at 14 DAG seedlings. n=10 replicates. E, Average length of B73 and *ehsp90.6* taproots at 14 DAG seedlings. n=10 replicates. Error bars = 10 average values ± SDs. ***, P< 0.001 (Student’s t test).

**Figure S2.**
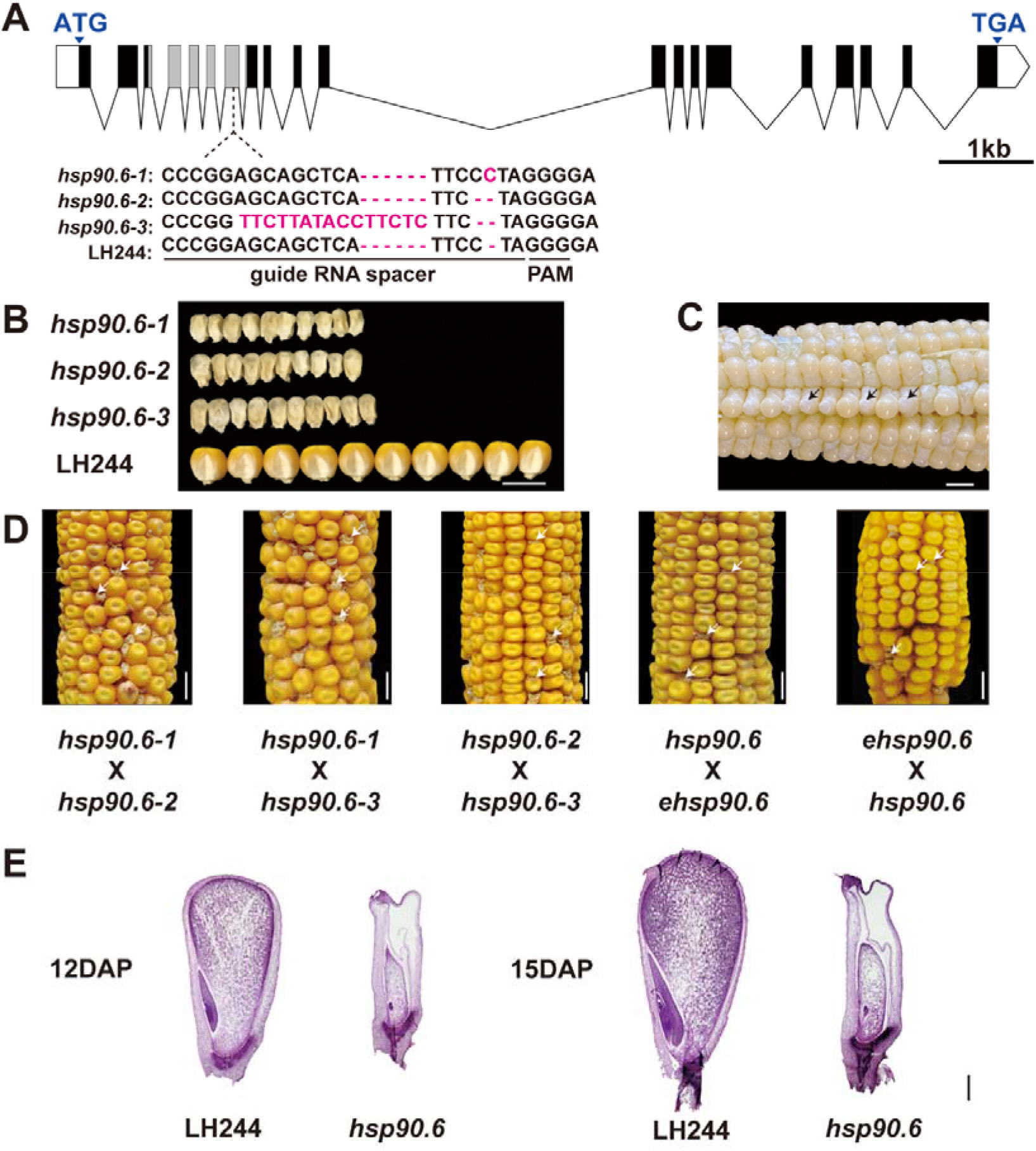
Phenotypic tests of ears of the CRISPR/Cas9-generated mutant *hsp90.6* A, CRISPR/Cas9-edited sequences of *hsp90.6* alleles. The gRNA spacer and protospacer-adjacent motif (PAM) site are indicated. B, Mature seeds of *hsp90.6* and LH244. Scale bar =1 cm. C, Ears of heterozygous *hsp90.6* mutants at 10 DAP. Scale bar =1 cm. D, Mature ears of heterozygous *hsp90.6* mutants crossed with different transformants events. Scale bar =1 cm. E, LH244 and *hsp90.6* seeds were observed at 12 DAP and 15 DAP under an optical microscope. Scale bar =1 mm.

**Figure S3.**
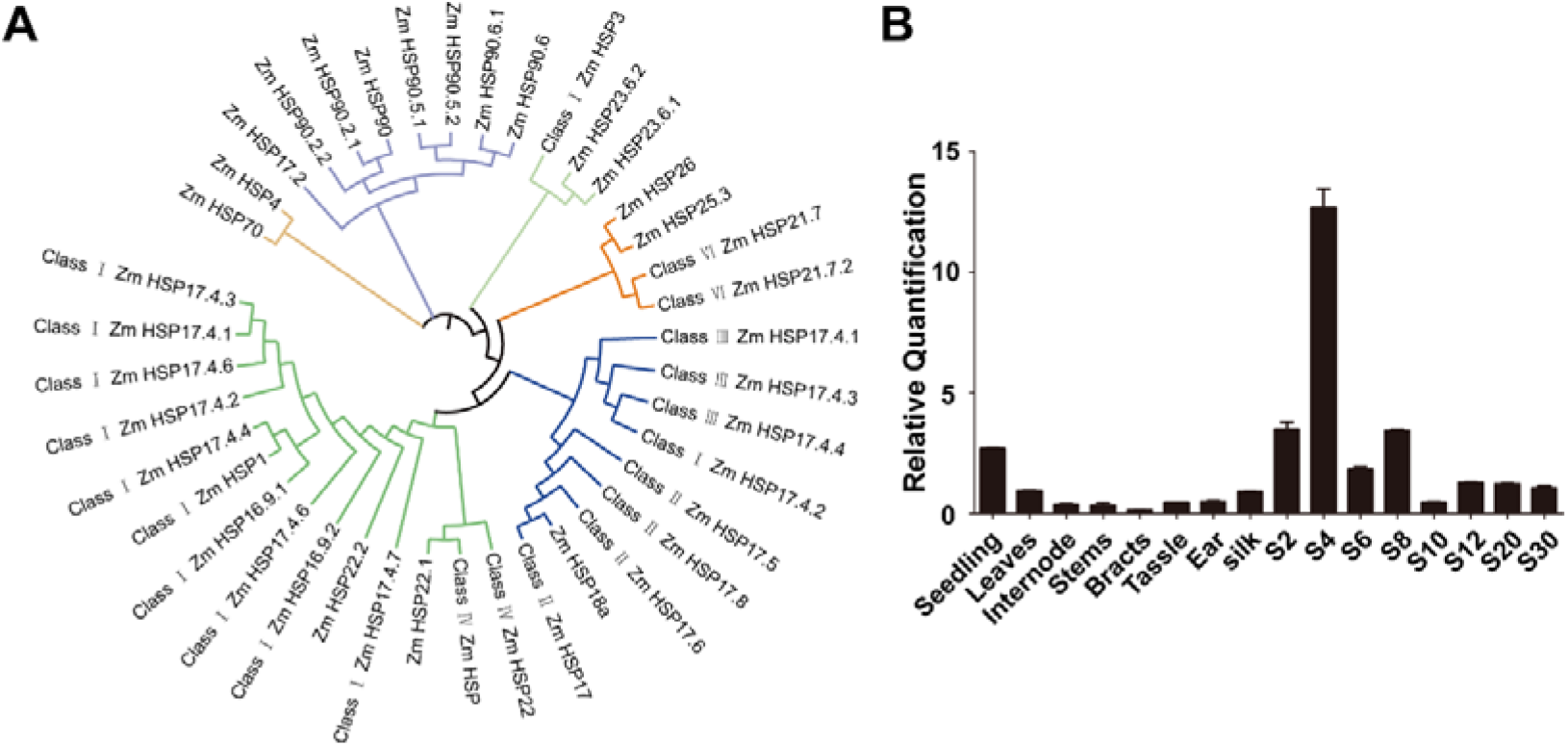
Characteristics of *HSP90.6* A, Phylogenetic relationships of the HSP family members in maize are known. B, RT–qPCR analysis of the expression of *HSP90.6* in different maize tissues. The transcript levels of *HSP90.6* were normalized to those of actin. For each sample, three independent biological replicates were analyzed. n = 3 replicates. Error bars = 3 average values ± SDs.

**Figure S4.**
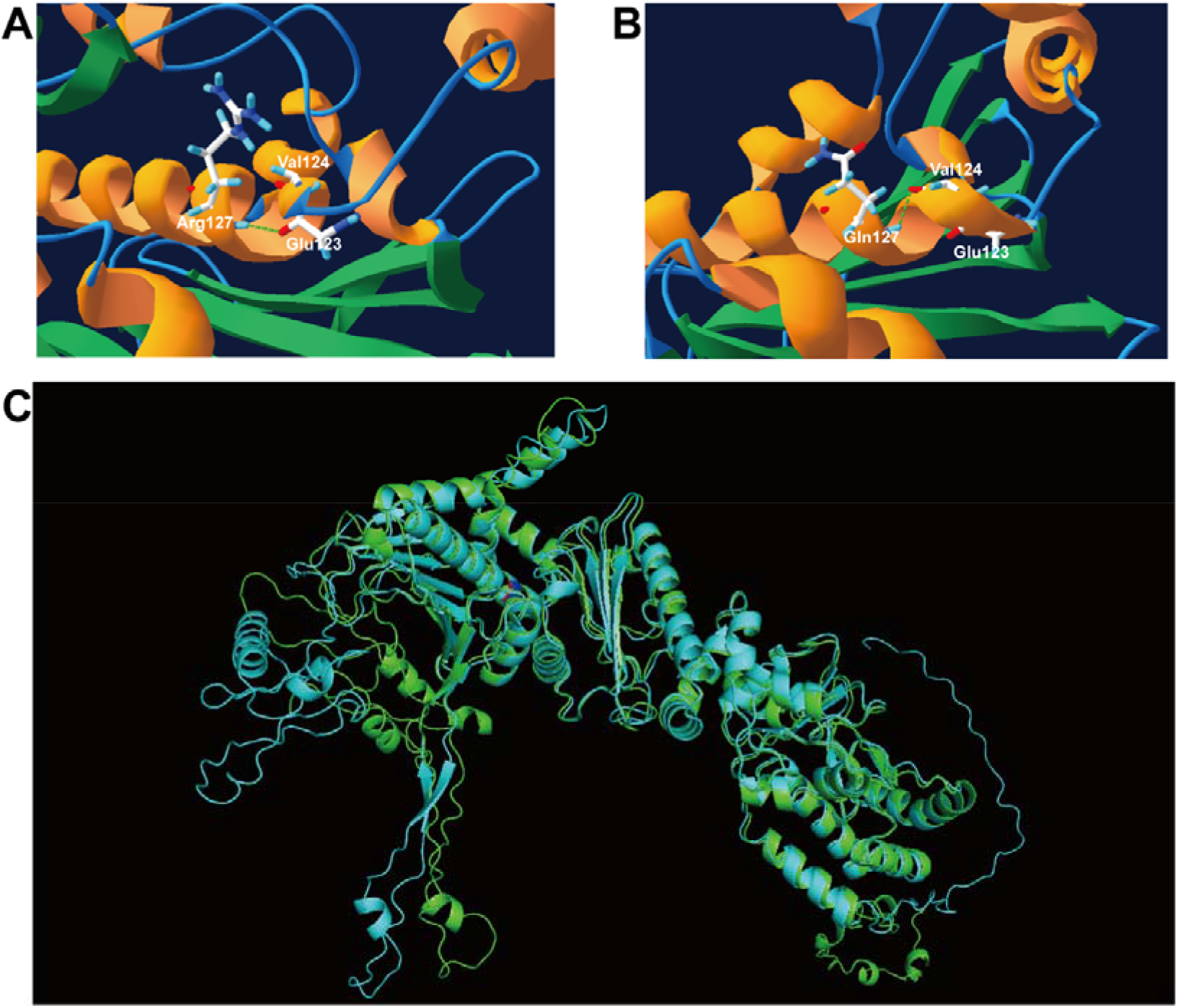
Prediction and comparison of protein structures A, B, Prediction of hydrogen bond distribution at the mutation site of WT (A) and *ehsp90.6* plants (B). C, Comparison of predicted protein structures comparison of WT and *ehsp90.6*. Green represents HSP90.6, cyan represents *hsp90.6*, and red and blue represent amino acid mutation sites.

**Figure S7.**
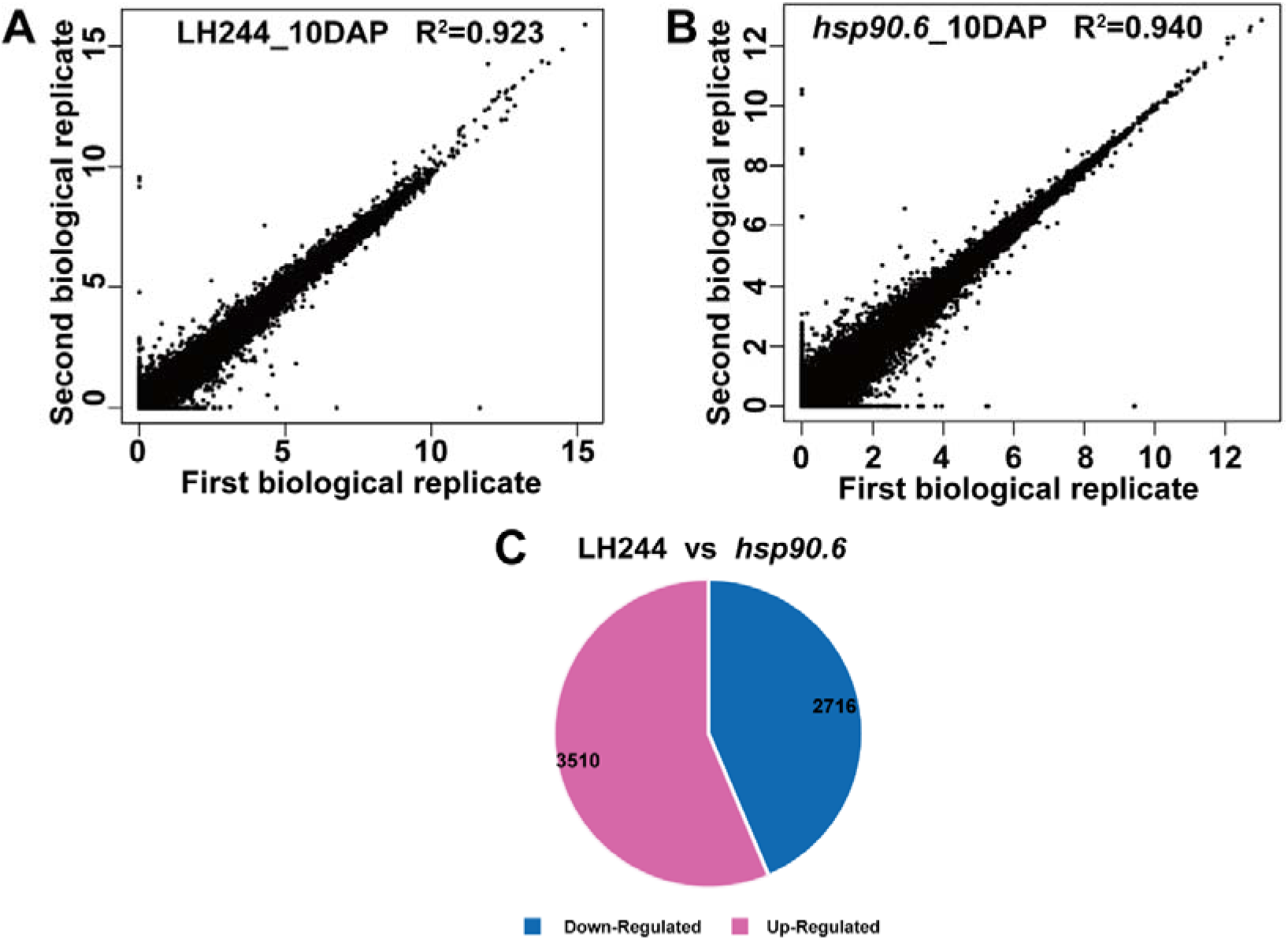
Analysis of DEGs A, B, Correlations between RNA-seq data of WT and *hsp90.6* (two biological replicates each). C, Genes differentially expressed between *hsp90.6* and the WT. (log2(fold-change)|>=1, Q value<0.05). The data are the means of two biological replicates.

**Figure S8.**
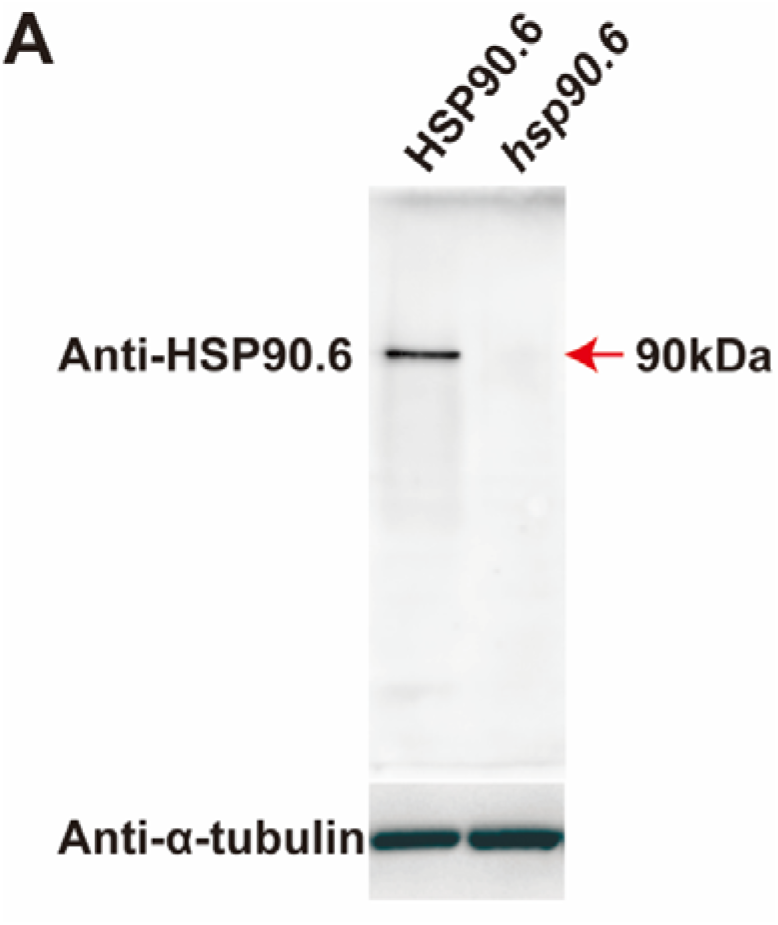
Western blot analysis of the specificity of anti-HSP90.6. The embryos and endosperm of WT and *hsp90.6* were isolated from ears at 10 DAP, and total protein was extracted for Western blotting. The protein levels were normalized to those of anti-α-tubulin.

**Figure S10.**
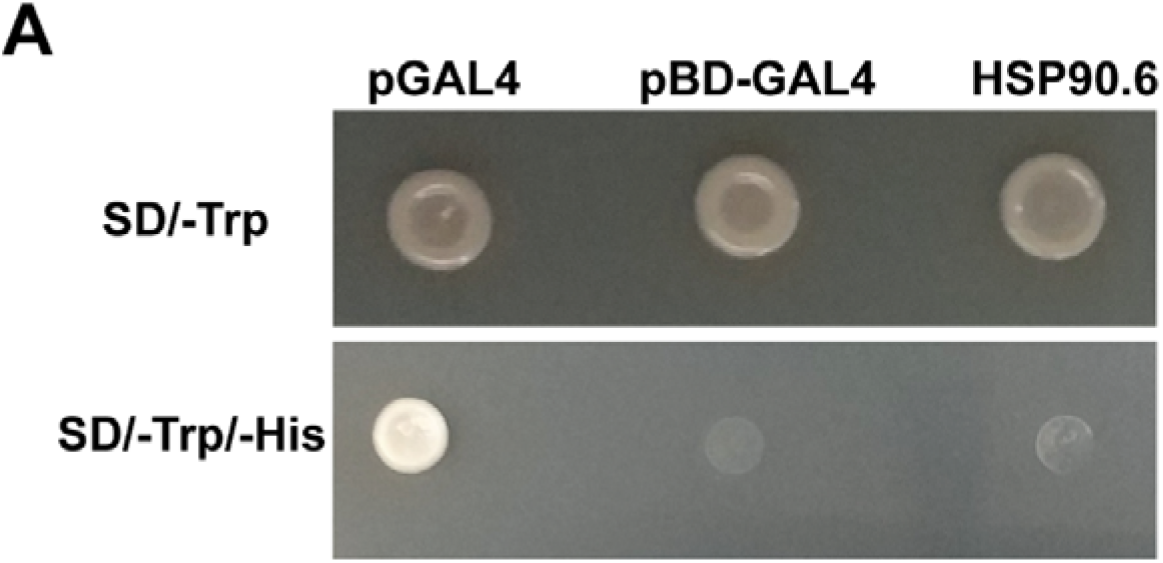
Analysis of transcriptional activation activity of HSP90.6. pGAL4, positive control; pBD-GAL4, negative control.

